# Video-based motion-resilient reconstruction of 3D position for fNIRS and EEG head mounted probes

**DOI:** 10.1101/621615

**Authors:** Sagi Jaffe-Dax, Amit H. Bermano, Yotam Erel, Lauren L. Emberson

## Abstract

**Significance:** We propose a novel video-based, motion-resilient, and fast method for estimating the position of optodes on the scalp.

**Aim:** Measuring the exact placement of probes (e.g., electrodes, optodes) on a participant’s head is a notoriously difficult step in acquiring neuroimaging data from methods which rely on scalp recordings (e.g., EEG and fNIRS), and is particularly difficult for any clinical or developmental population. Existing methods of head measurements require the participant to remain still for a lengthy period of time, are laborious, and require extensive training. Therefore, a fast and motion-resilient method is required for estimating the scalp location of probes.

**Approach:** We propose an innovative video-based method for estimating the probes’ positions relative to the participant’s head, which is fast, motion-resilient, and automatic. Our method builds on capitalizing the advantages, and understanding the limitations, of cutting-edge computer vision and machine learning tool. We validate our method on 10 adult subjects and provide proof of feasibility with infant subjects.

**Results:** We show that our method is both reliable and valid compared to existing state-of-the-art methods by estimating probe positions in a single measurement, and by tracking their translation and consistency across sessions. Finally, we show that our automatic method is able to estimate the position of probes on an infant head without lengthy offline procedures, a task which is considered challenging until now.

**Conclusions:** Our proposed method allows, for the first time, the use of automated spatial co-registration methods on developmental and clinical populations, where lengthy, motion-sensitive measurement methods routinely fail.

## Introduction

Functional near infrared spectroscopy (fNIRS) and electroencephalography (EEG) require knowledge of the positioning of probes on subjects’ scalp. Knowing the position of probes is an essential pre-requisite for key analytic steps: Aggregating results from a group of subjects, comparing between groups of participants (particularly from different developmental populations where head size systematically varies with group), and for estimating the source of the recorded signal (in fNIRS^1,2^). Desired positions of probes are determined *a priori* in accordance with either the standard positioning system (e.g., 10-20 in EEG^3^), or a predefined array that target specific cortical regions (typically in fNIRS^4^). However, it is not possible to place the probes exactly on the desired scalp locations for every experimental session. This is due to various reasons, including variability in head size and shape between participants, different practices between researchers, and even head growth between sessions in longitudinal studies with infants^5^. Therefore, there would be substantial improvement in analytic methods if there was a reliable, quick, and robust method for calibrating the 3D positions of the probes relative to the subject’s head.

Existing methods for 3D estimation of probes’ location are not suitable for early developmental or clinical populations and reduce the portability of these methods (one of their major benefits over MRI). Using the most popular approach, the 3D digitizer (e.g., Polhemus’ Fastrak or Patriot, Colchester, VT), to measure even a small number of points on the participant’s scalp typically requires a few minutes while a participant remains completely motionless^6^. Even with highly skilled developmental cognitive neuroscientists, early developmental populations (and many clinical populations) cannot meet this requirement and, thus, there are no published studies using a 3D digitizer for probe location estimation in early developmental populations (for older ages and limitations see refs ^7,8^). Furthermore, measurements with a 3D digitizer are often confounded by interference from metal objects in the participant’s surroundings. Thus, even in populations that can comply with these methods, it is difficult to move this method between experimental contexts. This is problematic as one of the major benefits of scalp-based neuroimaging is its portability (e.g., recording in rural Africa^9^). The 3D digitizer itself can cause interference with sensitive medical devices, such as cochlear implants, which make it non-suitable for this key clinical population.

Alternative co-registration methods have been used in developmental populations with some success but each has major limitations. Recently proposed photogrammetry methods also rely on tightly restricted movements of the subject and allow only a specific movement trajectory of the camera around the subject^14,15^. These methods require a lengthy recording of the participant (typically longer than two minutes; e.g., Delscan’s Artec or GeoScan^16^) or impose motionless requirements in an extremely specific environment (e.g., Philips’ GPS). Faster, more developmentally-friendly, co-registration methods are based on manual photogrammetry. These manual methods require expertise in image annotation^2,17^, are extremely time-intensive, and are prone to inter-researcher variability and biases. In these manual photogrammetry methods, several photos of the infant wearing the fNIRS cap are taken from a few different angles. The photos are then manually scaled according to a known size marker on the cap. The researcher then measures a few of the physical distances in the photos (e.g., nasion to inion, right ear to left ear, head circumference) and estimates the distances of the fNIRS channels from predefined fiducials^2^.

Semi-automatic co-registration methods require extensive physical measurements (i.e., using a measuring tape^10^) or MR scans of each subject wearing the fNIRS cap in the scanner^11–13^. These methods are also not suitable for developmental studies, for the same reasons specified above, involving motion restrictions for a few minutes and unpleasant environments. Other semi-manual photogrammetry methods use commercial 3D scanners, but still require manual annotation of probe locations on the output images^18^. Despite their disadvantages, these photogrammetric methods are currently being used for developmental studies^19,20^, since they offer the only reliable co-registration method in these populations. Our proposed video-based method takes the basic principles of these manual and semi-manual methods and extends it to an automatic video-based one.

Another existing method involves positioning probes on the scalp relative to skull landmarks (e.g., EEG positions^21,22^). These EEG positions can be then linked to the underlying cortical areas using existing atlases^23^. This method has the benefit of not requiring offline annotation as well as having the ability to locate central probes (i.e., the ones localized to EEG locations) across infants and populations. However, there are major limitations of this method. First, this method still requires minutes-long of measurements with the participant, rendering it challenging for infant studies. Second, given the fixed sensor distances of fNIRS (compared to EEG), even if one or a handful of probes are positioned carefully using skull landmarks, other probes are likely to vary across populations based on head size and shape. In other words, if a central probe position is positioned at Cz, the adjacent probes have a fixed configuration that typically is not changed based on head size or shape. Thus, this method (in combination with anatomical atlases) only allows alignment for a subset of probes.

In this paper, we present a novel video-based method that is both easy to implement by novice experimenters and is robust to participant’s head movements. Our method requires only ~20 seconds of video using widely available photographic equipment, recorded around the participant’s head, with the probes already mounted on the scalp. During acquisition, the participant can move his or her head freely without jeopardizing the accuracy of the measurement. We report both the validity and the reliability of our video-based method compared to the traditional 3D digitizer on a group of adult participants, and show it is as reliable as the 3D digitizer golden standard. Moreover, we also demonstrate the feasibility of this approach with early developmental populations – to which the 3D digitizer cannot be applied.

## Methods

Our automatic video-based method is depicted in Figure 1, and includes a preparatory step and four subject-specific steps. In a pre-processing step, we capture the cap in perfect conditions – well aligned on a plastic head. We then manually mark the positions of all fiducials and probes on the reconstructed 3D model, to create our reference *(model cap;* Figure 1A; see below fiducials’ definition). The model cap is used to learn the spatial relationship between fiducial points and probe positions. This allows us calculating the latter through interpolation of the former. This preregistration step is required for each new configuration of probe positions (i.e., for new regions of interest, caps, or populations). Then, for each subject we do the following: First, the participant’s head, with mounted cap and probes, is captured through a short video (Figure 1C), using an off-the-shelf mid to high-end camera (e.g., GoPro Hero6; GoPro, San Mateo, CA, see Figure 1B). Then, using a Convolutional Neural Network (Figure 1D), we crop away the background of each frame in the video, leaving only the cap and a part of the subject’s head (Figure 1E). This facilitates handling head motion, since it focuses the next steps solely on the cap, completely eliminating distractions. Next, we reconstruct a 3D model of the head using computer vision techniques for object 3D reconstruction (Structure from Motion – SfM^24^; Figure 1F). Lastly, we extract the coordinates of specific fiducials - an intuitive identification task once we already have a reconstructed model, in order to find all probe positions in MNI coordinate (Figure 1G).

**Figure 1.**
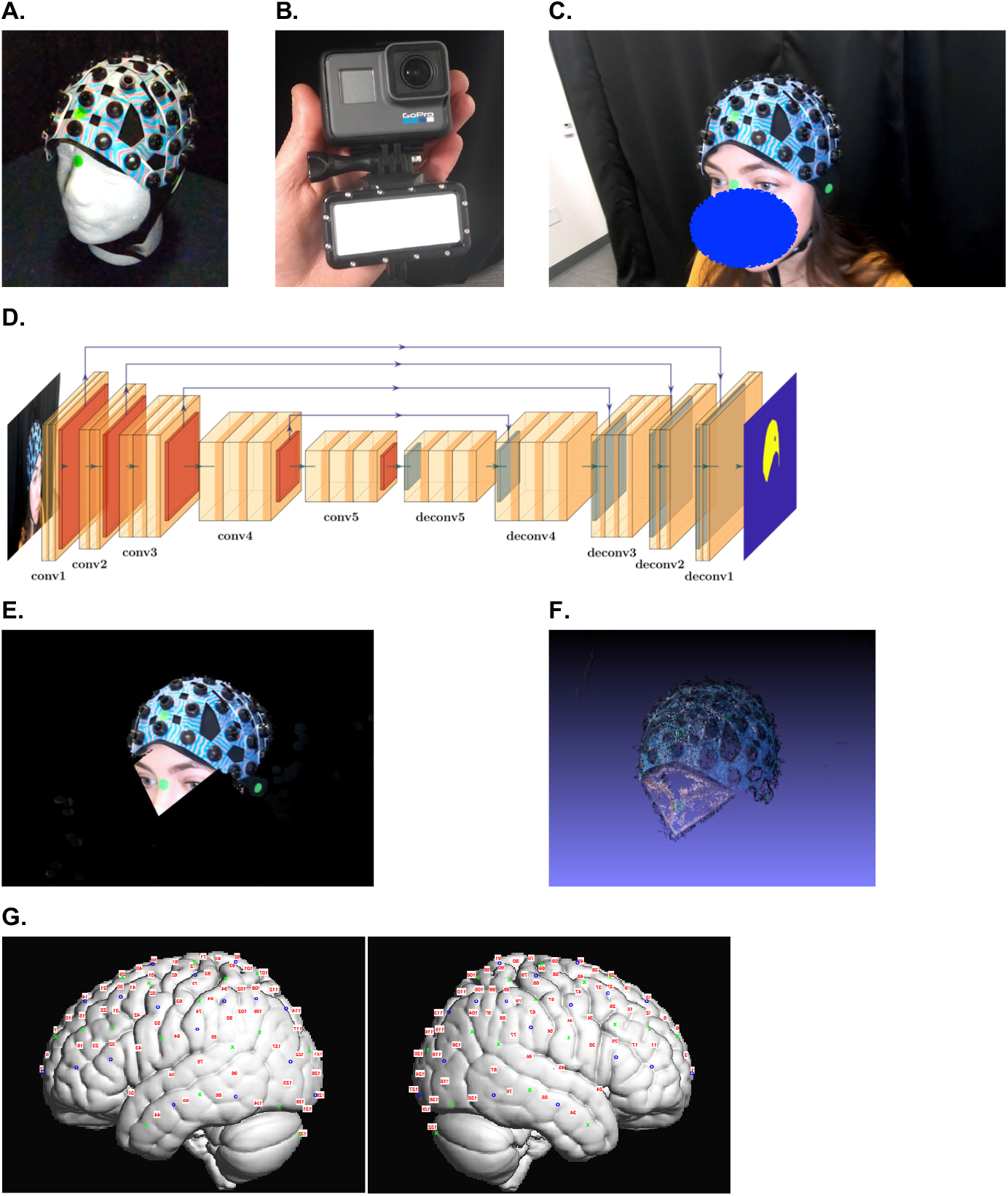
Method steps overview. A. The Model Cap is captured in perfect conditions, against a solid background with no movements. **B.** The short video is captured using an off-the-shelf camera with diffused flashlight. **C.** fNIRS cap is assembled with two-color pattern sheets and solid-color stickers placed in the fiducial positions, to be estimated. **D.** The architecture of the CNN used to create masks for the images. **E.** The frames are cropped using the mask from the neural net, a facial landmarks detector, and a sticker classifier. **F.** All cropped frames are combined to a 3D model through the popular Structure from Motion (SfM) computer vision technique. **G.** Fiducials are extracted from the model and channel positions are projected onto MNI space using SPM-fNIRS. Sources in blue circles; detectors in green crosses; channels in red numbers.

### Cap preparation

The basic requirement of the method is good tracking of the cap-mounted head throughout the SfM process, which is facilitated by feature-rich captured objects. To enrich the cap-mounted head with features, we replace the solid-color connector sheets of the fNIRS cap with multiple two-color patterned plastic sheets (red and blue in the example in Fig. 1B). We use Perlin noise^25^ to generate distinct patterns. A script for generating this specific pattern is in the Supplementary Materials, but any distinct non-trivial color pattern will suffice, e.g. drawings or cartoons. In addition, we place six solid color stickers (green in Fig. 1B) at specific positions on the cap: around the vertex, around the inion, between the vertex and the inion, and on the left, right, and front edges of the cap. These positions roughly correspond to Cz, Iz, Pz, T7, T8, and Fpz of the standard 10-20 system, respectively.

### Participant preparation

We fit the cap on the subject and place three additional solid color stickers (green in Fig. 1B) at specific anatomical positions on the participant fiducials: nasion (Nz), left preauricular (LPA), and right preauricular (RPA). In order to ensure other objects in the scene do not confuse our cropping method, we clear the surrounding of the participants from objects with similar color to the two-color plastic sheets and fiducial markers (e.g., no toys with similar colors, if the parent’s shirt had similar colors, we cover it with a dark barber cape, etc.). When the environment cannot be cleared of these object (e.g., infant clothing or pacifier), we allow the experimenter to choose the color of the solid-color stickers to be distinct from the subject’s clothing, and manually mark the chosen color after the fact (see next steps).

With this visual information about the spatial positions of the cap relative to the participant’s head, our problem boils down to finding the spatial relation between the points on the head to the those on the cap. Finding this relation is sufficient for exact positioning of the cap on the participant’s head.

### Video capture

As discussed below, the core of our method is based on finding the 3D positions of the sticker-marked fiducials using a computer vision algorithm. Since the 3D relationships between the fiducials is required, it is imperative all of them are clearly visible in the captured video, and that the computer vision system’s tracking is not lost during video acquisition. Along with patient comfort, these are the driving concerns for the process of capturing the video. Hence, we suggest the following procedure to help ensure successful tracking of each fiducial: We record the cap placement on the subject’s head using a GoPro Hero 6 Black. We use the slow-motion (240 fps) feature to reduce motion blurring. Of course, any camera with a similar slow-motion option will suffice for the requirements of our method (e.g., iPhone 8 or higher). Furthermore, we attach a diffused flashlight to the camera to compensate for uneven lighting conditions across all head sides (see Fig. 1B). Lastly, we start and end the video recording at approximately the same position relative to the subject’s head. Note that depending on environmental conditions not all of, or even none of, the steps are necessary for success. These steps however, have proven to be reliable and simple to implement. An example for such a video recording is available at: https://youtu.be/sD75ctsGlD4.

### Cropping the video frames

Passing irrelevant information about the background of the given object-to-be-reconstructed to the SfM method often yields bad reconstructions, partial reconstructions or falsely merged objects. This undesired reconstruction results are especially common in the presence of motion, i.e., when the object is moving during the video acquisition. Head movement, and subsequent bad reconstruction, are the major challenges that existing automated photogrammetry co-registration methods face with early developmental populations.

In order to avoid passing irrelevant information about the background to the SfM stage and to prevent the issues that arise from participants’ movements, we crop everything but the head and cap from each video frame. First, we maintain color consistency as much as possible, through white-balance correction of the video frames using OpenCV^26^ package for Matlab^27^. Then, the user manually marks, from a single frame, the color of the stickers (green in Fig. 1) that were placed on the fiducial points. Manually selecting the color of the solid-color stickers allows flexibility regarding the choice of used stickers – one could determine a fixed color for the fiducial stickers, which would render the last aforementioned step unnecessary. However, the sticker color should be distinctly different from any other color in the scene (see participant preparation stage).

After these steps, we use the following automatic processes to produce a cropping mask to each frame:

1. Identify the pixels corresponding to the fiducial stickers: Each cluster of at least five connected pixels with a hue within ± 0.15% of the sticker hue (measured in HSV space) is classified as a sticker. A circle is defined around each cluster with an additional margin of 20% of the radius.
2. Identify the cap in each video frame: We designed and trained *CapNet* (available in Supplementary), a Convolutional Neural Network (CNN) based on the segmentation neural network^28^, pre-trained on the CamVid^29^ dataset and fine-tuned on our dataset. We found that manually annotating 100 images with pixel-wise classes of “cap” and ‘‘background” creates a sufficient training set. The convolutional layers used in the architecture composing the encoder part are ((width x height x depth) x stack): (360×480×64)x2, (180×240,128)x2, (90×120×256), (45×60×512)x3, (22×30,512)x3. The encoder is followed by a symmetrical stack of de-convolutional layers composing the decoder (Fig. 1C). Batch-Normalization layers are placed after each convolutional layer, followed by a ReLu activation layer. Max-Pooling layers are used between every stack of convolutional layers, and each one of them additionally has a skip connection to its Max-Unpooling counterpart. Images are resized to fit the input layer using bicubic interpolation. Network training was done using the stochastic gradient descent with momentum optimizer, with momentum value of 0.9, for 100 epochs. Network training stopped after reaching the very satisfactory global accuracy of 98%, and weighted average (across classes) intersection over union (IoU) of 90%.
3. Identify participants’ face: We found that keeping a part of the subject’s face in the image, in addition to the cap, improves the 3D reconstruction of the nasion region. The participant face is identified using an off-the-shelf neural-network-based facial detector, found in OpenCV^30^. We define a polygon connecting the nose, external eye-edges and forehead, and add it to the cap in the region of the frame that is kept for further analysis. The application of such an example polygon can be seen in Fig. 1E.

The rest of the frame is blackened (Fig. 1E), and frames in which more than 98% of the frame is blackened are rejected from further processing. These frames are either blurred frames due to fast motion of the camera, or frames in which the participant’s head is out of frame.

### 3D surface reconstruction

In order to find the 3D coordinates of the fiducial points, a 3D model is automatically reconstructed. The cropped frames are passed to a Structure from Motion^31^ software (Visual SfM^32^). This widely-used algorithm searches for shared features between frames (or *correspondences*^33^), and estimates the camera position for each frame accordingly. Note that understanding this mechanism had enabled our key head-motion robustness contribution - when searching for correspondences between frames, a static background along with a moving head confuses the camera-position-estimation process. Hence, by removing the background in the cropping step, a moving head can be simply interpreted as additional camera movement, rendering accurate and successful reconstructions of our object of interest. We instruct the software to only search for correspondences that are less than two seconds apart. Searching for temporally distant correspondences is very computationally expensive, and might yield faulty matches, since the video does not show the same parts for long periods. In addition, since the videos are taken such that they start and end the same way, matches are also searched between the first and last frames of the video. This closed-loop helps in reducing accumulated drift, and in cases where matches tracking is lost mid-way through the video. A 3D model can then be built (Fig. 1F), using the correspondences, camera positions, and simple geometric triangulations.

### Fiducial extraction and probe positions estimation

Our basic assumption in this part is that the cap undergoes limited deformation when worn by the participant (e.g., the cap does not stretch according to differences in head size as an EEG cap would). Instead, differences in head size, shape, cap placement, etc. are captured by the positions of the markers on the cap relative to the head (cap points and fiducial points, respectively, see below). After these points are extracted, we assume the cap can only rotate globally with respect to the head, and scale along the anterior-to-posterior and left-to-right axes. In this way, head size is implicitly considered in our reconstruction of probe positions. Therefore, capturing five points on the cap (along with the ones on the head) are sufficient for complete cap-head registration. The position of the probes can be then interpolated similarly to common photogrammetry-based methods^2^. In practice, we used 6 points for redundancy, with two in the difficult-to-capture back region (“Pz” and “Iz”), because infants are sitting on their parent’s lap.

From the reconstructed 3D model, we identify the position of the fiducials by clustering nearby reconstructed vertices, which hold the solid sticker color. Using RANSAC^34^, we determine the fiducial name associated with each aforementioned cluster (by looking at the expected relative position of all fiducials). In other words, we select three of the identified points, and assign a labeling to them, taken from the initial cap model. We then deform the head so that the three points match their counterparts and measure the distance of the rest of the markers from the ones they end up closest to. After choosing the labeling that minimizes the latter distances, we use the head fiducials (Nz, AR, AL, Cz) to project the cap points (“Cz”, “Pz”, “Iz”, Front, Left, Right of the cap; we are assuming Cz is relatively in the correct position) into MNI space by rotation and scaling^35^. The channel positions are then interpolated in MNI space based on the pre-registered model cap relative to the cap points using SPM for fNIRS (Tak et al., 2016; Fig. 1G). The interpolation is performed using barycentric coordinates: We represent each manually marked probe as a weighted sum of all fiducials *f_i_, i* = 1…9. In other words, we find all *w_i,j_* ’s such that 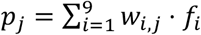 in the pre-registration step, and simply use the same formula with the newly found fiducials to extract probe positions.

### 3D digitizer measurement

For comparison to our automated video-based method of 3D reconstruction, we collected the 3D coordinates of the head points and cap points using a 3D digitizer (Fastrak, Polhemus, Colchester, VT) twice for each adult subject in each session. We then projected the channel positions into MNI space using the same pre-defined model cap and SPM for fNIRS.

### Estimation of validity and reliability of channel position estimation

We compared the channel positions (139 channels; LABNIRS, Shimadzu inc., Kyoto, Japan) determined based on our new video-based method to the positions found using the dominant method in adult participants, the 3D digitizer (Fastrak, Polhemus, Colchester, VT). We did the comparison twice for each subject (*N* = 10) for the same session and compared the positions of the same channels both between the two methods (inter-methods validity) and within each one (intra-method reliability). The intra-method comparison was used to estimate the error of each method independently (test-retest reliability).

In addition, we measured the channel position for a separate group of adult subjects (*N* = 10) in two separate sessions (separate days, multiple cap placements) to estimate whether the video-based method captured the shifts in cap positioning on the same subject as compared to the 3D digitizer.

### Feasibility for early developmental populations

Finally, we show how this method can be used to estimate probe positions with early developmental populations. Specifically, we present results from a 6-month old infant. The example video is available here: https://youtu.be/ecn-GSGoRh8. While it would be ideal to compare our video-based reconstruction methods as conducted in infants to other methods, we cannot conduct the same comparison as adults because it is not practically possible to collect the digitized positions of the fiducials and probes using the 3D digitizer. After months of effort on this front and communication with other developmental cognitive neuroscience labs, we reached the conclusion that this method is too sensitive to motion to be reliable and, thus, is unusable for this population.

## Results

### Video-based method, validated through the standard 3D digitizer

We found that the validity of our new video-based method was highly comparable to the field’s standard method – the 3D digitizer (Fig 2A-B right). Namely, the distances between the positions of the same channel as estimated by the two methods were similar, albeit slightly larger, to the distances between the positions of the same channel as measured by the 3D digitizer twice (inter-methods validity of 3.4 ± 0.9 mm compared to intra-method reliability of 2.6 ± 0.6 mm; Mean ± STD; *t*_9_ = 3.1, *P* < 0.05). Thus, there is a high correspondence between the channel positions estimated by our video-based methods and the 3D digitizer.

**Figure 2.**
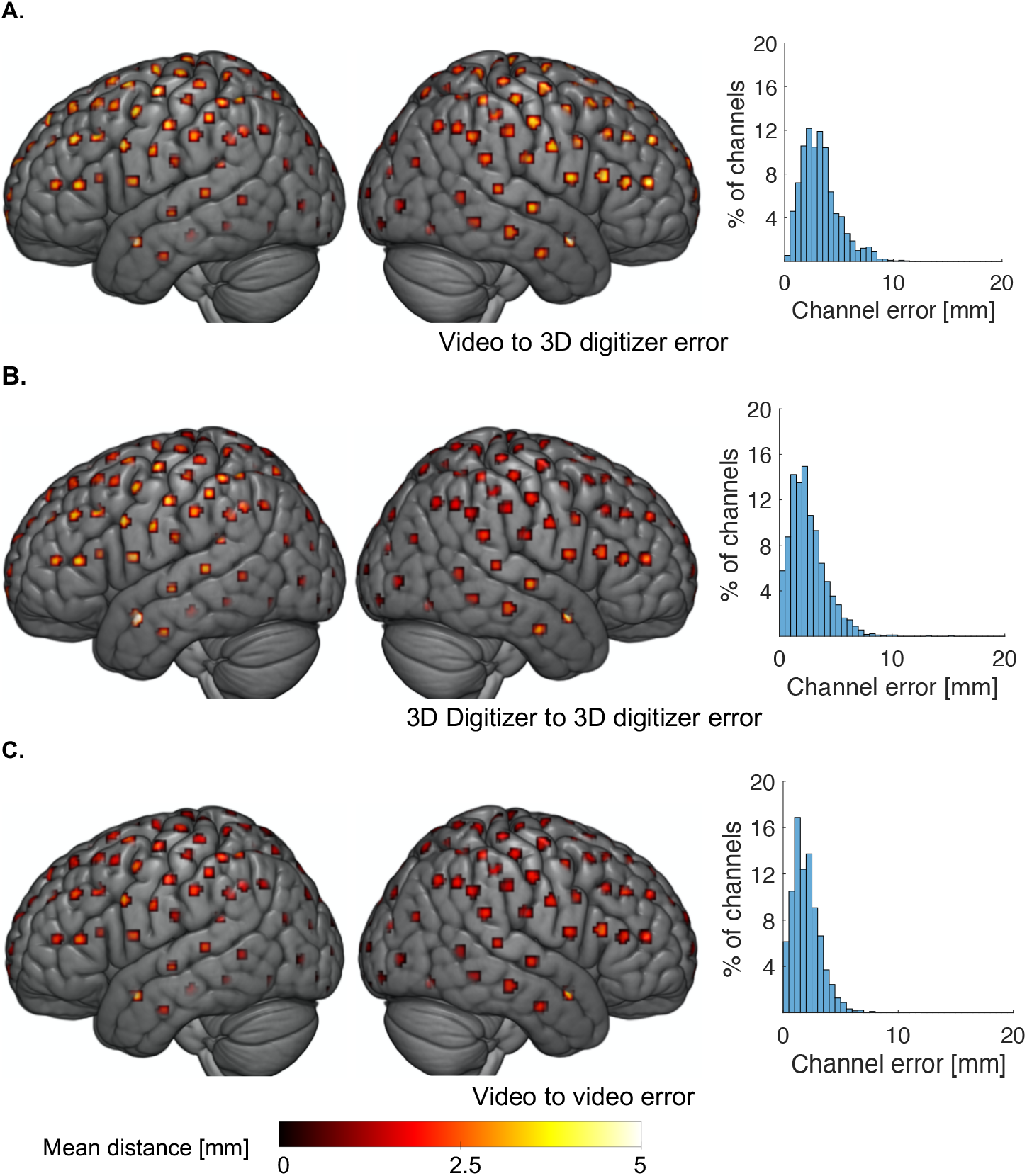
Spatial distribution of inter-method validity and intra-method reliability, comparing our method and the digitizer. **A.** Distance from our (video) method to the 3D digitizer one (inter-method validation; 3.4 ± 0.9 mm). **B.** Distance between two measurements of the 3D digitizer (intra-method reliability; 2.6 ± 0.6 mm). **C.** Distance between two measurements of video-based method (intra-method reliability; 2.0 ± 0.5 mm). Distances are represented by the color and the diameter of the patches for each channel position.

Moreover, the reliability of our video-based method was better than the reliability of the 3D digitizer. The distances between the estimated positions of the same channel as measured twice by our video-based method (2.0 ± 0.5 mm; Fig 2C right) was smaller than that of the 3D digitizer (*t*_9_ = 2.6, *P* < 0.05; paired t-test). Overall, the magnitude of the errors that we found is much smaller than the radius of the channels, whose positions were estimated (typically on the order of centimeters).

While the discrepancies between the two position estimation methods were spread evenly through the scalp, we did observe a tendency for larger distances in the anterior temporal channels (Fig. 2A left and middle). This increased error at this position stems from smaller reliability of both methods in these areas (Fig. 2B-C left and middle). The lower reliability of both methods in the anterior temporal lobe is probably a result of the low angle of camera or digitizer that is required to acquire the position of the fiducials in this region.

### Video-based method captures cap positioning variability

We now considered how well these two localization methods identify any change in position of the cap (i.e., shift or displacement of each channel) for the same participant across sessions. We found a significant regression coefficient for the size of the estimated shift in channel position (in mm) between the two methods (Red line in Fig. 3A; *F*_1,1388_ = 5.86 *p* < 0.05; Linear mixed effect model). Namely, the shift size as estimated by the video-base method predicts the shift size as estimated by the 3D digitizer method. As Figure 3A clearly shows, there were many cases where the two estimation methods did not agree on the shift size between the two sessions (e.g., a high shift size was estimated by the 3D digitizer and a small shift size was estimated by the video-based method for the points on the righthand side; channels below the diagonal – black dashed line in Fig. 3A). We manually examined a few cases of large discrepancies between the shift estimation of the two methods. In most cases, the video-based method was actually closer to estimating the shift magnitude correctly. For example, in Figure 3C, the shift between the sessions of channel 66 (circled in yellow) for participant 1 was estimated as 3.6 mm by the video-based method and 24.5 mm by the 3D digitizer. Our manual estimation, based on image-space measurements scaled by known distances between channels, suggests a shift of 7mm – a number much closer to the video-based estimation than to the 3D digitizer-based one (Fig. 3C).

**Figure 3.**
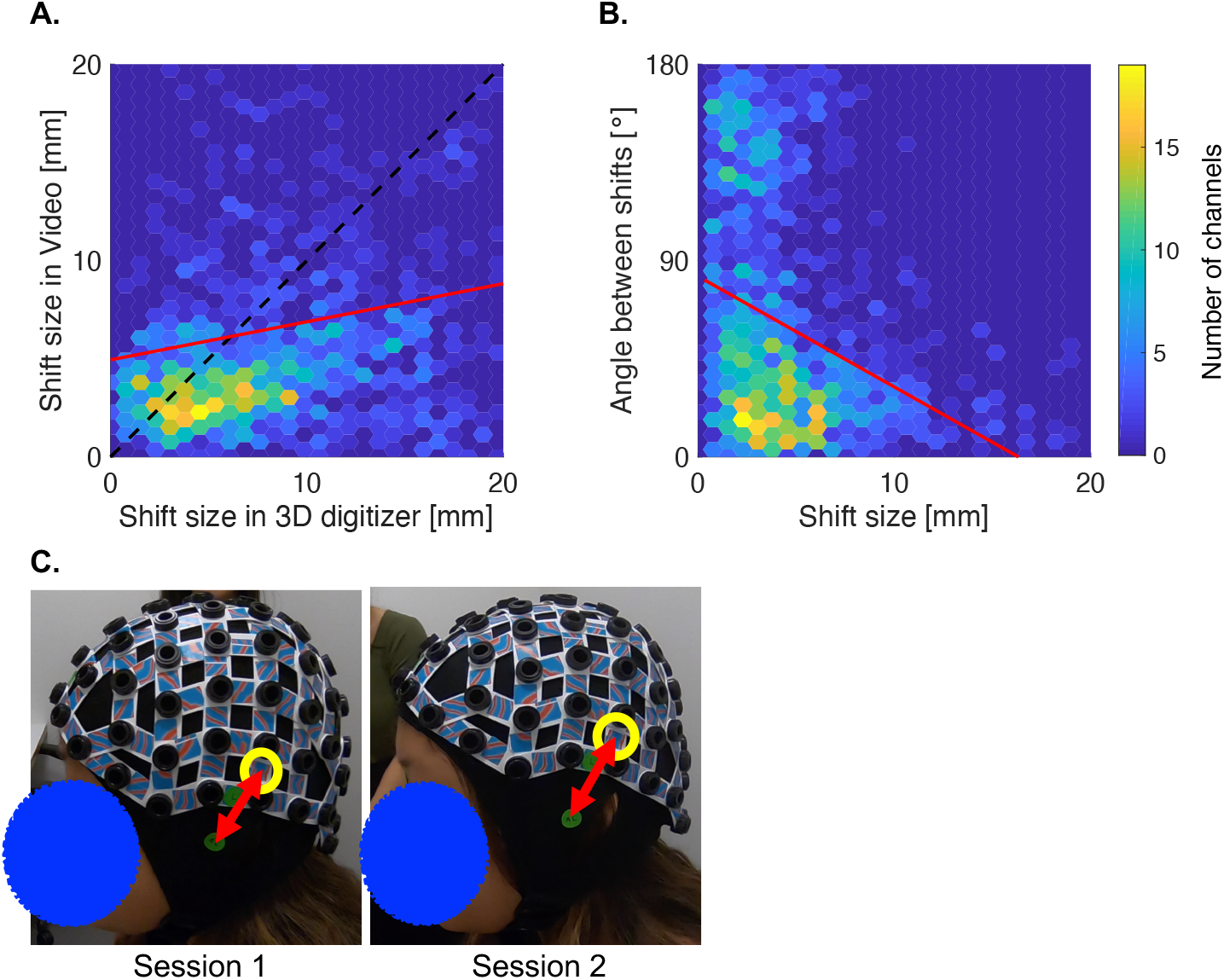
Comparison of estimation of cap shifting between session through the video-based and the 3D digitizer methods. **A.** Estimation of shift size between the sessions of the video-based method vs the 3D digitizer one. The color of each hexagon represents the number of channels in its range. We found a high correlation between the shifts as estimated by the two methods (*F*1,1368 = 21.8 *p* < 10^-5^; red line). **B.** Angles between the shift as estimated by the video-based method and the shift as estimated by the 3D digitizer method vs the size of the shift between sessions. The distribution of angles in skewed towards zero *(K* = 0.3, *p* < 10^-25^; distribution along the ordinate). We found a negative correlation between the size of the angle between estimated shifts and the size of these shifts. When the estimated shifts are larger, the direction of the shifts, as estimated by the two methods were similar (smaller angle). **C.** Manual estimation of a case of large discrepancy between the two estimation methods. Left - photo of participant 1 in session 1. The distance of channel 66 (circled in yellow) from the left tragus (lower green sticker) was 49.3 mm (the length of the red arrow). Right - photo of participant 1 in session 2. The distance between channel 66 and the left tragus was 53.3 mm. The shift of the channel position between the two sessions was ~4 mm and was closer to the video-based estimation (3.6 mm) than to the digitizer-based estimation (24.5 mm).

We also found that the shifts in channel positions were estimated in similar directions using the two methods. We measured the spatial angle between the direction of the shifts as estimated by the video method and the direction of the shift as estimated by the 3D digitizer. The estimated shift of each channel between sessions can be represented as a line between the two 3D estimated positions. Each method thus estimates such a line and we can measure the level of agreement between the methods by measuring the angle between these lines. The angle between the direction of the shifts, as estimated by the two methods, was small (closer to zero than a uniform distribution; distribution along the ordinate in Fig. 3B; *K* = 0.3, *p* < 10^-25^; Kolmogorov-Smirnov test; Mean angle ± STD: 60.6° ± 50.1°). Crucially, the similarity in shift direction was found to be greater in the larger shifts (Red line in Fig. 3A; *F*_1,1368_ = 21.8 *p* < 10^-5^; Linear mixed effect model), where this similarity in direction matters most (i.e., when the cap shifts substantially and may potentially result in a change in the cortical region that the channel is assigned to).

Investigating these shifts in cap placement across sessions reveals how crucial it is for any co-registration to be accurate. Even if recordings of the same subject take place with the same cap mere days apart, these shifts in cap placement are large enough to change the cortical region under a channel. In our sample, more than 8% of the channels were shifted to a different lobe between sessions.

Additionally, our automatic method does not rely on the specific head size, or models it specifically. Instead, it automatically adjusts for the head size, by measuring the distance between the fiducial points and the cap directly, taking the head size implicitly into account (see *Fiducial extraction and probe positions estimation* section of the Methods). Thus, we expect no correlation between the error magnitude and head sizes. Unfortunately, we did not take independent measurements of our subjects’ head sizes (e.g., circumference, inion to nasion distance, etc.). However, we qualitatively compare the errors that were obtained for a distinctly larger head with the errors that were obtained for a smaller one. We did not find any systematic difference (Fig. 4). We also find in the visual comparison between these two head sizes that the probes are shifted down on the smaller head and up on the larger head reflecting the differences in the relative size of the cap to the participant’s head size. This logical transformation reflects the fact that head size is accommodated by our method through differences in the fiducial points and cap points and doesn’t need to be included explicitly.

**Figure 4.**
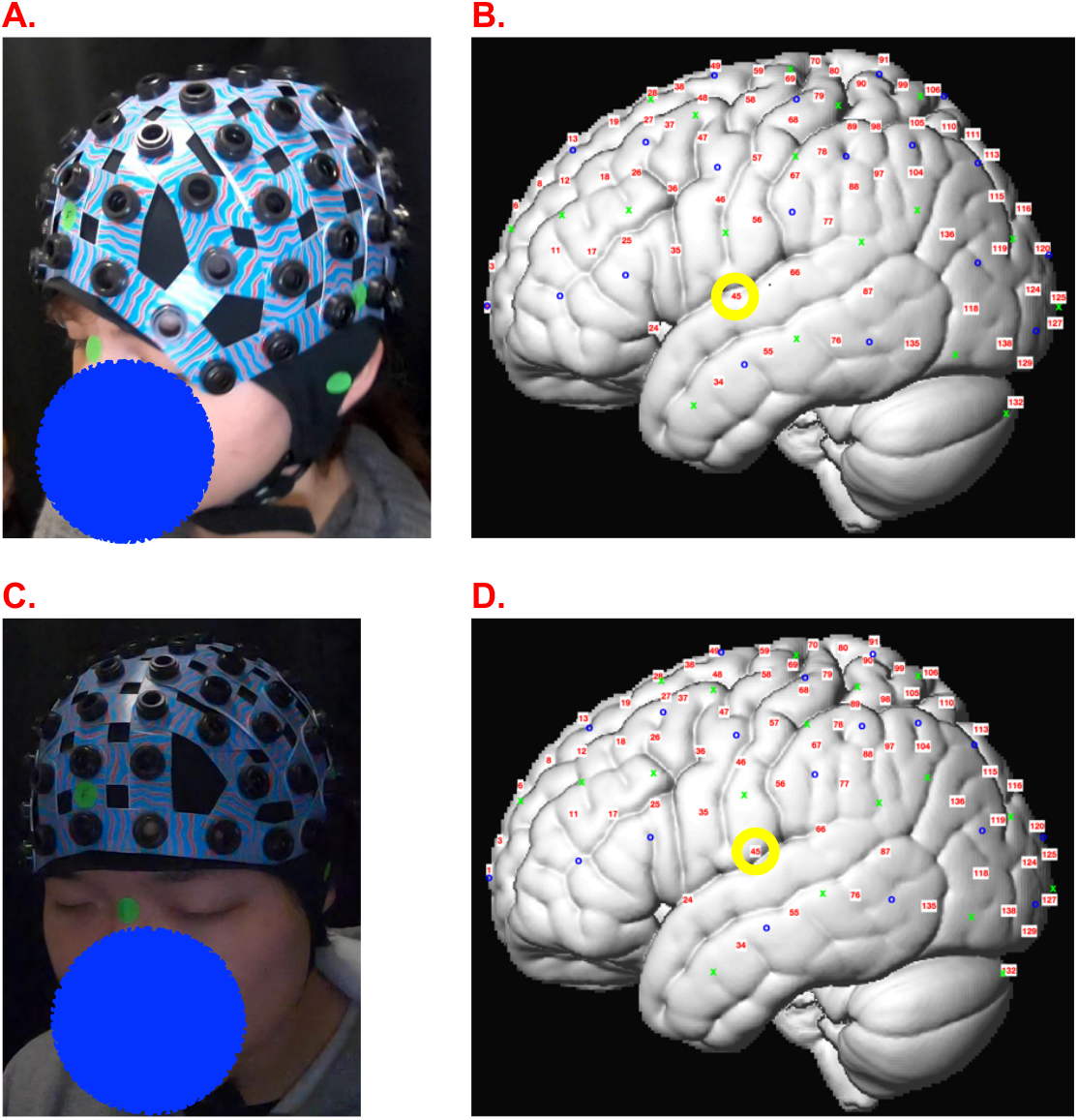
Comparison of two subjects with relatively distinct head sizes. The subject with the smaller head (A) had a mean error of 2.7 mm. The subject with the larger head (C) had a mean error of 2.3 mm. Due to the rigidity of the fNIRS cap, the probe positions on the smaller head (C) were shifted slightly downwards relative to the larger head (D). For example, channel 45 (circled in yellow) was estimated in STG on the smaller head size but in post-central gyrus on the larger head size.

### Automatic estimation of channel position in infants

We successfully reconstructed a 3D model from video recordings of infant participants and estimated the channel positions on their scalp through our automated video-based method (using methods described above, Fig. 6 for one sample infant). Thus, here we present a proof of concept that the same method can be used for early developmental populations. In practice, we have employed this method for many dozens of infants at different ages. Since there can be no comparison with a 3D digitizer, we focus this paper of the validity of the overall method in adult participants in comparison to established methods.

As apparent in this example, the cap was slightly misplaced on the infant, tilting to the right (Fig. 5A). Our probe position estimation method successfully estimated this misplacement (Fig. 5D).

**Figure 5.**
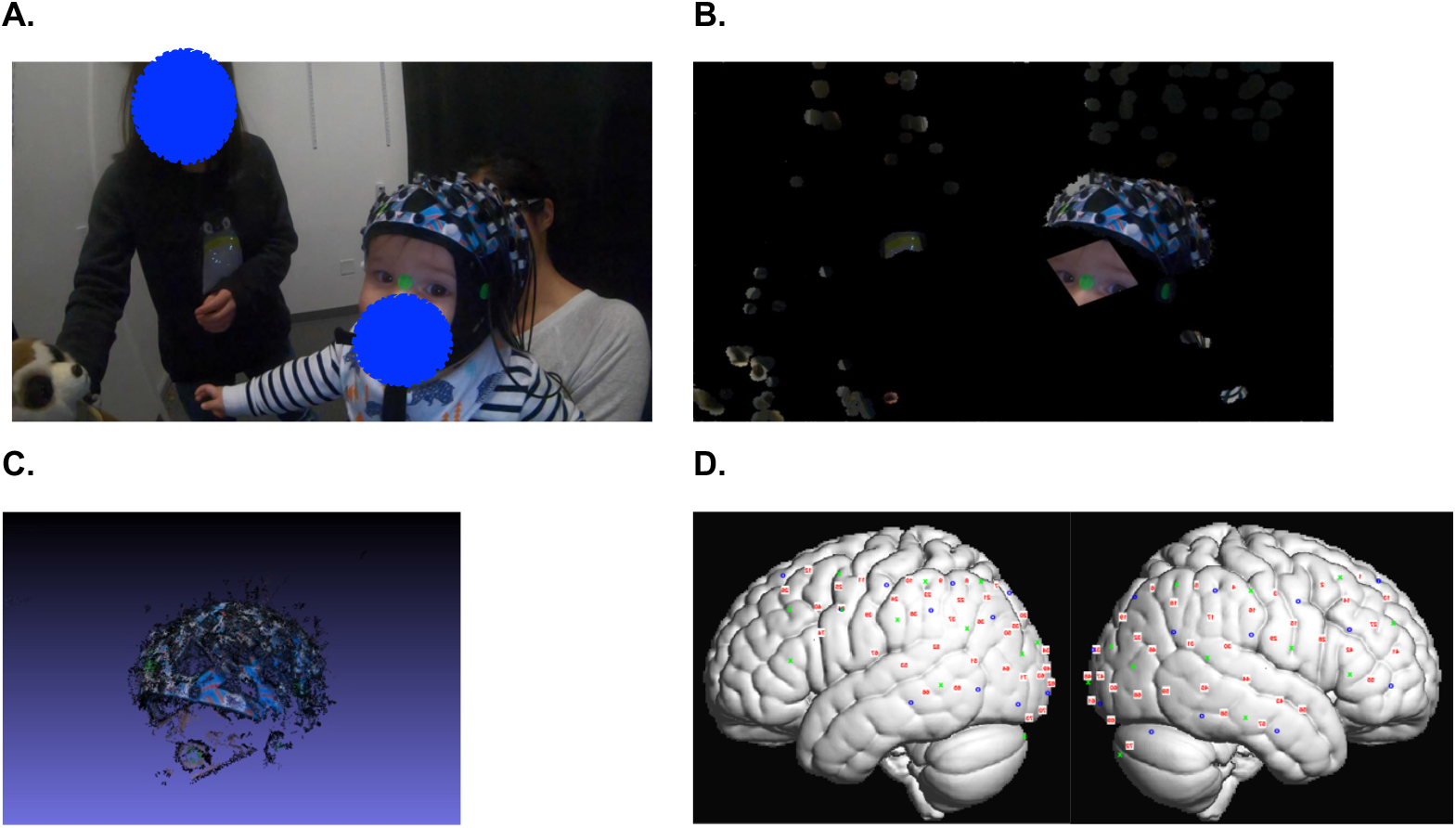
Infant channel position estimation using the video-based method. **A.** The captured video. **B.** The images were cropped to eliminate confounds of movements. **C.** A 3D model was reconstructed using Visual Structure from Motion. Coordinates of fiducial points were extracted from the model. **D.** Channel positions were interpolated from fiducial coordinates onto MNI space.

## Discussion

The field of developmental cognitive neuroscience requires an automated reliable method to estimate the spatial position of probes on the scalp. This method is especially important when researchers are drawing conclusions about the development of functional anatomy that underlies the recorded signal from EEG or fNIRS. As we show in this work, even minor unintentional shifts in cap positioning can yield dramatic changes between the cortical region that is being recorded in different sessions. Such uncertainty in position estimation and changes between sessions may lead to increased levels of noise in neuroimaging results and even to false conclusions regarding the underlying cortical activity.

We present an innovative approach for 3D localization of probes relative to a participant’s head, which is highly needed in scalp-based methods of neuroimaging such as EEG and fNIRS and particularly for developmental and clinical populations where current methods are inadequate. The proposed method requires only a short video recording during the experimental session. Moreover, during this video, participants can move freely and interact with research staff and caregivers. These differences in methodology make this method suitable for developmental and clinical populations.

We designed this new method to substantially improve the ability to conduct probe localization in developmental and clinical populations. Currently available methods have major disadvantages: Some are susceptible to participant’s movements and take a relatively long amount of time^10^. Thus, they are not useable for developmental populations (3D digitizer) and are onerous for clinical populations^37^. Other methods require laborious and bias-prone manual annotation (manual co-registration^2^). The video-based method we present here is both motion-resilient and automatic. Additionally, this method does not require constant power supply and is not sensitive to metal in the environment. Therefore, this method is applicable not only to the population it was designed for, but also to out-of-lab settings and may be widely used in clinical settings and global neuroscience projects. The automatic analysis of the videos does not require the extensive manual annotation of images and thus is less prone to bias and much less laborious.

We present evidence that the proposed method is both valid and reliable compared to the standard method currently employed in the field, the 3D digitizer. Overall, we find strong agreement between these methods of localization. We find a high level of agreement between methods (inter-method validity), and within the method in a test-retest repeated measurements configuration (intra-method reliability). Importantly, our video-based method exhibits better intra-method reliability compared to the traditional 3D digitizer. In the region where these two methods had the lowest agreement, the anterior temporal region, we suspect that it stemmed from difficulty in placing the digitizer in the correct angle, perpendicular to the scalp. Throughout the scalp, the error size of our estimation is an order of magnitude smaller that the size of the channels that are to-be-estimated.

We further tested the localization abilities of these methods by examining changes in localization of channels between sessions with the same subject. We find that these methods similarly estimate the displacement of the cap across sessions. We estimated the session-to-session variability in cap placement using the video-based and the digitizer methods and found good agreement between the two. As can be expected, we found that the agreement between the two estimation methods about the shift direction was higher when the size of the shift was larger. Where there were disagreements between the methods (e.g., large difference between the size of the estimated shift), we have found that the video localization method was usually more accurate in channel position estimation.

Perhaps most importantly, we have demonstrated the successful measurement of infants’ head mounted probe positions: An estimation that is not possible using currently available online methods (e.g., 3D digitizers). Of course, successfully estimating misalignments in cap placement is of even greater value when administering experiments with developmental population, since it is a more prevalent problem in such cases.

Using a different model cap, as described at the beginning of the method section, researchers can easily adapt the outline proposed here to their own system and to any cap type.

The method of course has some limitations. First, much like other methods, it is acutely dependent of recovering all the sticker-marked fiducials (excluding the one redundant pair). Unlike other methods though, the missing information can be recovered from the abundance of visual information found in the captured video. Secondly, while quick to capture, the method takes a few minutes to compute the registration results, meaning the technician is not aware of the quality of the registration while running the experiment. Therefore, future implementation of the proposed video-based method will include a user interface, facilitating easier insertions of new cap models, as mentioned above, and other features such as manual corrections to badly captured or detected positions. This approach potentially can profit from automation wherever possible but from human understanding where it is not. Another interesting direction to take would be to develop a single neural-network based pipeline producing a real-time indication of the calibration results. This could be done perhaps by collecting online feedback from users of the system regarding specific reconstructions.

Finally, we hope our proposed co-registration method will allow the combination of data from different locations and labs and will enable the creation of a unified anatomical framework for fNIRS analysis.

## Acknowledgements

This research was supported by grants from the National Institutes of Health (R00 4R00HD076166-02), McDonnell Foundation (220020505), and the Bill and Melinda Gates Foundation – Modelling Neurodevelopment: Physical Growth above the Neck. All authors declare that they have no conflict of interests.

